# Long-term dietary exposure to a mixture of phthalates enhances estrogen and beta-catenin signaling pathways, leading to endometrial hyperplasia in mice

**DOI:** 10.1101/2024.09.16.613339

**Authors:** Ritwik Shukla, Athilakshmi Kannan, Mary J. Laws, Amy Wagoner Johnson, Jodi A. Flaws, Milan K. Bagchi, Indrani C. Bagchi

**Affiliations:** Departments of Comparative Biosciences, University of Illinois at Urbana-Champaign, Urbana, IL; Molecular & Integrative Physiology, University of Illinois at Urbana-Champaign, Urbana, IL; Mechanical Science and Engineering, University of Illinois at Urbana-Champaign, Urbana, IL; Carle R. Woese Institute for Genomic Biology, University of Illinois at Urbana-Champaign, Urbana, IL; Carle Illinois College of Medicine, University of Illinois at Urbana-Champaign, Urbana, IL

## Abstract

Phthalates, synthetic chemicals widely utilized as plasticizers and stabilizers in various consumer products, present a significant concern due to their persistent presence in daily human life. While past research predominantly focused on individual phthalates, real-life human exposure typically encompasses complex mixtures of these compounds. The cumulative effects of prolonged exposure to phthalate mixtures on uterine health remain poorly understood. To address this knowledge gap, we conducted studies utilizing adult female mice exposed to a phthalate mixture for 6 and 12 months through ad libitum chow consumption. We previously reported that continuous exposure to this phthalate mixture for 6 months led to uterine fibrosis. In this study, we show that the exposure, when continued beyond 6 months to 1 year, caused fibrotic uteri to display hyperplasia with a significant increase in gland to stroma ratio. Endometrial hyperplasia is commonly caused by unopposed estrogen action, which promotes increased expression of pro-inflammatory cytokines and chemokines and proliferation of the endometrial epithelial cells. Indeed, RNA sequencing analysis revealed a marked upregulation of several estrogen-regulated genes, Wnt ligands that are involved in oncogenic pathways, as well as chemokines, in phthalate-exposed uterine tissues. Consequently, the exposed uteri exhibited increased proliferation of endometrial epithelial cells, and a heightened inflammatory response indicated by extensive homing of macrophages. Further studies revealed a marked enhancement of the Wnt/β-Catenin signaling pathway, potentially contributing to the development of endometrial hyperplasia. Collectively, this study underscores the significance of understanding the exposure to environmental factors in the pathogenesis of endometrial disorders.

## INTRODUCTION

Phthalates, which are esters of phthalic acid, are synthetic chemicals extensively utilized as plasticizers in a variety of consumer products (1–5). High molecular weight phthalates with long chains include di(2- ethylhexyl) phthalate (DEHP), di-iso-nonyl phthalate (DiNP), di-iso-decyl phthalate (DiDP), di-n-octyl phthalate (DnOP), and di(2-propylheptyl) phthalate (DPhP). These are incorporated into polyvinyl chloride. Conversely, short-chain phthalates are employed in the production of personal care items such as perfumes, nail polish, deodorants, and lotions. This group comprises dimethyl phthalate (DMP), diethyl phthalate (DEP), benzyl butyl phthalate (BBzP), di-n-butyl phthalate (DnBP), and di-iso-butyl phthalate (DiBP). A significant quantity of phthalates is utilized globally each year (6). As phthalates are not covalently bound to plastics, they easily leach into the environment, resulting in continuous human exposure through inhalation, ingestion, and dermal contact (7). Ingestion is the most prevalent exposure pathway (8,9). Research reveals that phthalate metabolites are detected in nearly 100% of human urine samples tested (10–12). Notably, phthalate metabolite concentrations are higher in women than in men, likely due to the greater use of personal care products by women (13).

Growing scientific evidence associates phthalate exposure with adverse health effects due to the endocrine-disrupting nature of these chemicals. Epidemiological studies suggest that phthalate exposure correlates with reduced pregnancy rates, increased miscarriage rates, and preeclampsia in women (12,14–19). Additionally, phthalate exposure is linked to adverse pregnancy outcomes, such as low birth weight and compromised intellectual development and growth in children (15,17). Previous studies using rodents have shown that prenatal exposure to phthalates affects folliculogenesis, delays puberty, and reduces fertility (20,21). It has also been reported that exposure to DEHP results in uterine abnormalities in mice, including decreased proliferation of luminal epithelium and an increased number of abnormally dilated blood vessels in the endometrium (22). Previous studies demonstrated that while exposure to DiNP impairs the ability of mice to maintain pregnancy, mice exposed to DEHP did not experience pregnancy loss unless fed a high-fat diet (23–25). Although these studies have primarily focused on exposure to a single phthalate over a limited timeframe, humans encounter a mixture of phthalates daily throughout their lives.

Hence, we designed a study in which female mice were chronically exposed to a phthalate mixture via chow ad libitum and analyzed the effects of this exposure on the uterus. The composition of the phthalate mixture was DEHP, DiNP, BzBP, DBP, DiBP, and DEP, which was derived from the Illinois Kids Development Study (I-KIDS), where the levels of phthalate metabolites were determined in the urine samples from pregnant women (26). Long-term dietary exposure to this mixture adversely affects estrous cyclicity (27).

In our previous study, we reported that continuous exposure to this phthalate mixture for 6 months promotes uterine fibrosis (28). In this study, we report that exposure to the phthalate mixture for 1 year causes endometrial hyperplasia accompanied by enhanced estrogen-dependent signaling, inflammatory response, and activated Wnt/β-Catenin signaling pathway.

## MATERIALS & METHODS

### Chemicals

The phthalates, which were 98% pure, were obtained from Sigma-Aldrich (St. Louis, Missouri). Corn oil from Columbus Vegetable Oils (Des Plaines, Illinois) served as the vehicle control.

### Animals

Female CD-1 mice aged 33 days and male CD-1 mice aged 7 weeks were obtained from Charles River Laboratories (Wilmington, Massachusetts) and housed in the College of Veterinary Medicine vivarium at the University of Illinois Urbana-Champaign (Urbana, Illinois). As previously described, the female mice were group housed in polysulfone cages (Allentown, Allentown, New Jersey) at a density of 3 mice per cage, with 1/8 corn cob bedding (Shepherd Specialty Papers), environmental enrichment (iso-BLOX, catalog #6060, Envigo), and reverse osmosis purified water (27). All animal handling, housing, and procedures were approved by the University of Illinois Institutional Animal Care and Use Committee.

### Study design and dosing

Phthalates were administered to mice ad libitum in the rodent chow as previously described (27). The base chow was a modified version of the AIN-93G formulation (TD.94045), where soybean oil was replaced with corn oil from Columbus Vegetable Oils (Des Plaines, Illinois). Chow containing 7% corn oil served as the vehicle control group. Phthalates, mixed in corn oil, were provided to Envigo for chow preparation. The phthalate mixture used in this study consisted of 35% DEP, 21% DEHP, 15% DiNP, 15% DBP, 8% DiBP, and 5% BzBP, designed based on phthalate metabolite concentrations in the urine of pregnant women in central Illinois (26). High phthalate levels are typically found in food contact materials like plastic wrap and containers, making ingestion a significant exposure route for humans (9). To replicate human exposure, the phthalate mixture was administered via rodent chow at doses of 0.15 ppm and 1.5 ppm. Each treatment group comprised 12–14 mice. Based on previous studies, the doses were selected assuming that a 25 g mouse consumes approximately 5 g of food per day (29–31). Therefore, the 0.15 ppm dose equates to about 24 mg phthalate/kg body weight/day, and the 1.5 ppm dose to about 200 mg phthalate/kg body weight/day. These doses fall within the range of daily human, infant, and occupational exposure (32,33). Beginning at 6 weeks of age, the mice were continuously treated with the chow for 12 months. This continuous daily exposure to phthalates closely mimics human exposure patterns.

### Tissue collection and analysis

At 6 months and 1 year of age, the mice were euthanized during diestrus, and their uterine horns were collected. One portion of the uterine horn was fixed in 10% neutral buffered formalin (NBF), while the other portion was flash frozen in liquid nitrogen for subsequent gene expression analysis. For histological examination, the fixed uteri were dehydrated through a graded series of ethanol concentrations (from 70% to 100%). The samples were then embedded in paraffin, sectioned, and stained with hematoxylin and eosin.

### Immunohistochemistry (IHC)

Uterine tissues were processed and subjected to immunohistochemistry (IHC) as described previously (25). Briefly, paraffin-embedded tissues were sectioned at 5 μm and mounted on microscopic slides. Sections were deparaffinized in xylene, rehydrated through a series of ethanol washes, and rinsed in water. Antigen retrieval was performed by immersing the slides in 0.1M citrate buffer solution, pH 6.0, followed by microwave heating for 25 min. The slides were allowed to cool and endogenous peroxidase activity was blocked by incubating sections in 0.3% hydrogen peroxide in methanol for 15 min at room temperature. After washing with PBS for 15 min, the slides were incubated in a blocking solution for 1 h. This was followed by incubation overnight at 4°C with antibodies specific for KI67 and CTNNB1. Pictures were taken using the Olympus BX51 microscope equipped for fluorescent imaging and connected to a Jenoptik ProgRes C14 digital camera with c-mount interface containing a 1.4 Megapixel CCD sensor. Fluorescent images were processed and merged using Adobe Photoshop Extended CS6 (Adobe Systems).

#### Quantitative Real time PCR analysis (qPCR)

For gene expression analysis, RNA was converted to cDNA, and real-time quantitative PCR was performed using SYBR-green master mix (Applied Biosystems) on a QuantStudioTM 3 Real-time PCR instrument (Applied Biosystems). The mean threshold cycle (Ct) for each sample was calculated from Ct values obtained from three replicates. The normalized ΔCt in each sample was determined by subtracting the mean Ct of the reference gene from the mean Ct of the target gene. ΔΔCt was then calculated as the difference between the ΔCt values of the control and treated samples. The fold change in gene expression relative to the control was calculated using the 2−ΔΔCt method for relative quantification. The mean fold induction and standard error of the mean (SEM) were calculated from at least three independent experiments.

### Statistical Analysis

Statistical analyses were conducted following methods previously established (34). Data were presented as mean ± SEM. Statistical tests included a two-tailed Student’s t-test and Mann-Whitney rank sum test for single comparisons, one-way analysis of variance (ANOVA) with Bonferroni post-test for multiple comparisons between samples or time points, and two-way ANOVA with Bonferroni post-test for multiple comparisons across different samples and time points. Additionally, an equal variances analysis was performed on all numerical data to determine the appropriateness of parametric or non-parametric hypothesis tests. Statistical significance was set at P ≤ 0.05 and is indicated by asterisks in the figures. All data were analyzed and plotted using GraphPad Prism 9.4 (GraphPad Software).

## RESULTS

To investigate the effects of long-term dietary phthalate exposure on the uterus, adult mice were fed chow containing a vehicle control (corn oil) or a mixture of phthalates at doses 0.15 ppm and 1.5 ppm, as previously described (27). Uterine sections obtained from control and mice exposed to the phthalate mixture for 1 year were analyzed by hematoxylin and eosin (H&E) staining to study changes in uterine histology. Interestingly, we observed a dramatic increase in the number of uterine glands in the 0.15 ppm treatment group compared to controls (Fig. 1, middle panel). We found that exposure to the higher dose, 1.5 ppm, did not show a pronounced effect on the glands to stroma ratio compared to the 0.15 ppm exposure level. Thus, to gain insights into the mechanisms underlying the development of glandular hyperplasia in the uterus, we focused on the 0.15 exposure level.

**Figure 1:**
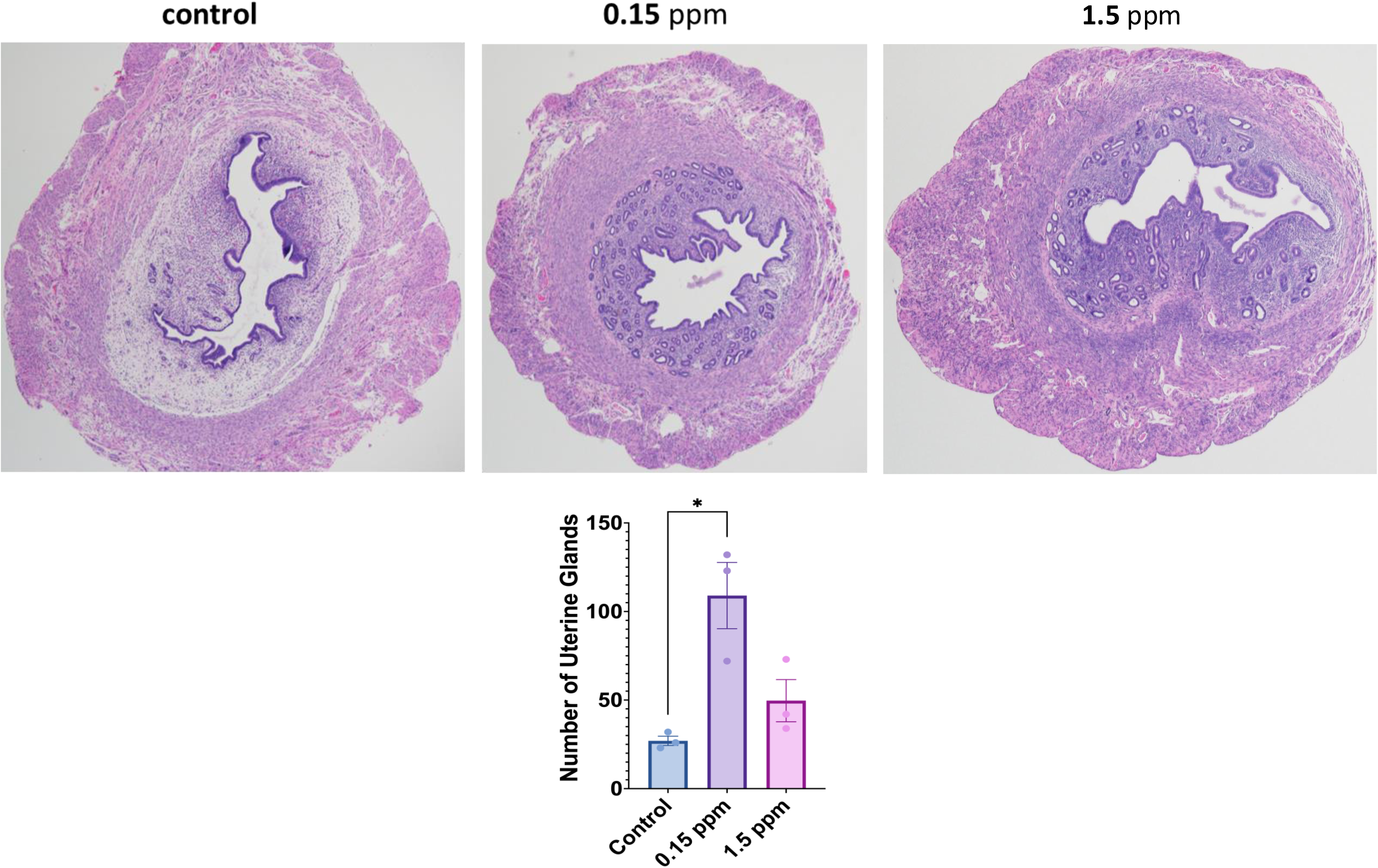
Histological examination of uterine sections. Upper: Serial sections of uterine horns of control (a), 0.15 ppm (b), and 1.5 ppm (c) of phthalate mixture exposed mice at 12 months were examined by H&E staining. The representative images from the control and phthalate mixture treated groups are shown. **Lower:** The number of glands in the uterus was determined at random locations. Data represent mean ± SEM from three separate samples. Asterisks indicate statistically significant differences (*P < 0.05).

To obtain unbiased insights into biological processes impacted by phthalate exposure, we performed RNA-sequencing to compare the expression levels of transcripts in the uteri of control and mice exposed to a 0.15 ppm phthalate mixture for 6 months (28). Our study revealed an upregulation of several estrogen-regulated transcripts in the phthalate-exposed uteri compared to the untreated controls (35,36). The RNA-seq data were validated by real-time PCR analysis (Fig. 2A). We confirmed a marked upregulation of mRNA levels corresponding to *Cx43*, *Muc1*, *Hif2α*, *Fgf9, Ccn1*, and *Six1* in uterine samples of 0.15 ppm groups indicating that a long-term exposure to the phthalate mixture enhances estrogen signaling (35–40). Interestingly, many upregulated genes, such as *Fgf9*, *Ccn1*, *Pbx3*, *Six1,* and *Wt1,* are known to be involved in oncogenic pathways. We also observed increased expression of a subset of Wnt ligands, such as *Wnt4*, *Wnt9a*, *Wnt5b*, *Wnt7a,* and *Wnt8b,* upon exposure to the phthalate mixture. These Wnt ligands are tightly associated with cancer, including Wnt4, known to be regulated by estrogen signaling in the uterus (Fig. 2B). Besides these factors, we also observed an upregulation of CC chemokine ligands (CCL), *Ccl2*, *Ccl17*, *Ccl2*, *Ccl21a*, *Ccl7,* and *Ccl8* (Fig. 2C). These are a subset of cytokines known to be involved in inflammation. Collectively, these results indicated that long-term exposure to the phthalate mixture enhances estrogen-dependent signaling, which is closely linked with cell proliferation and inflammation.

**Figure 2:**
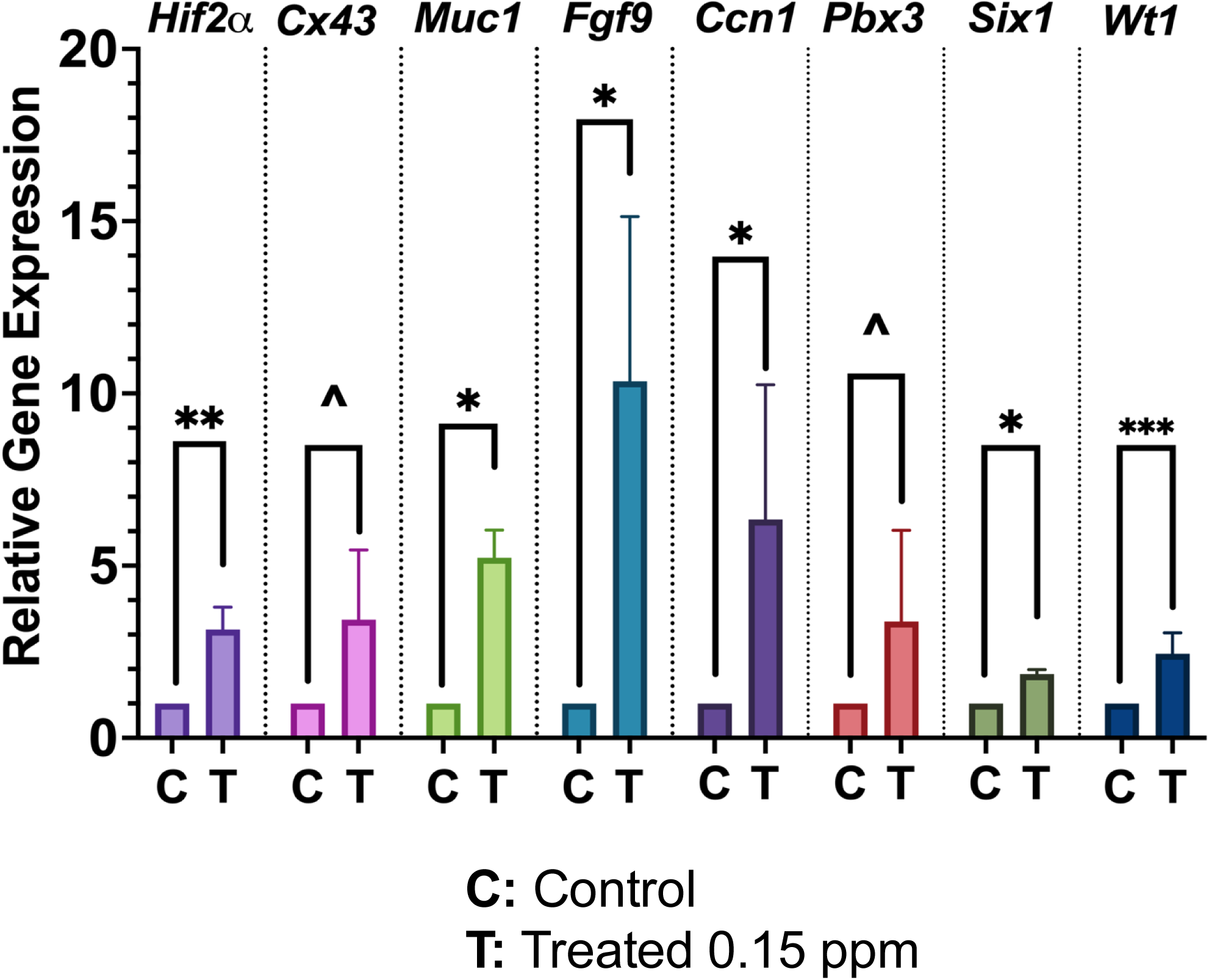
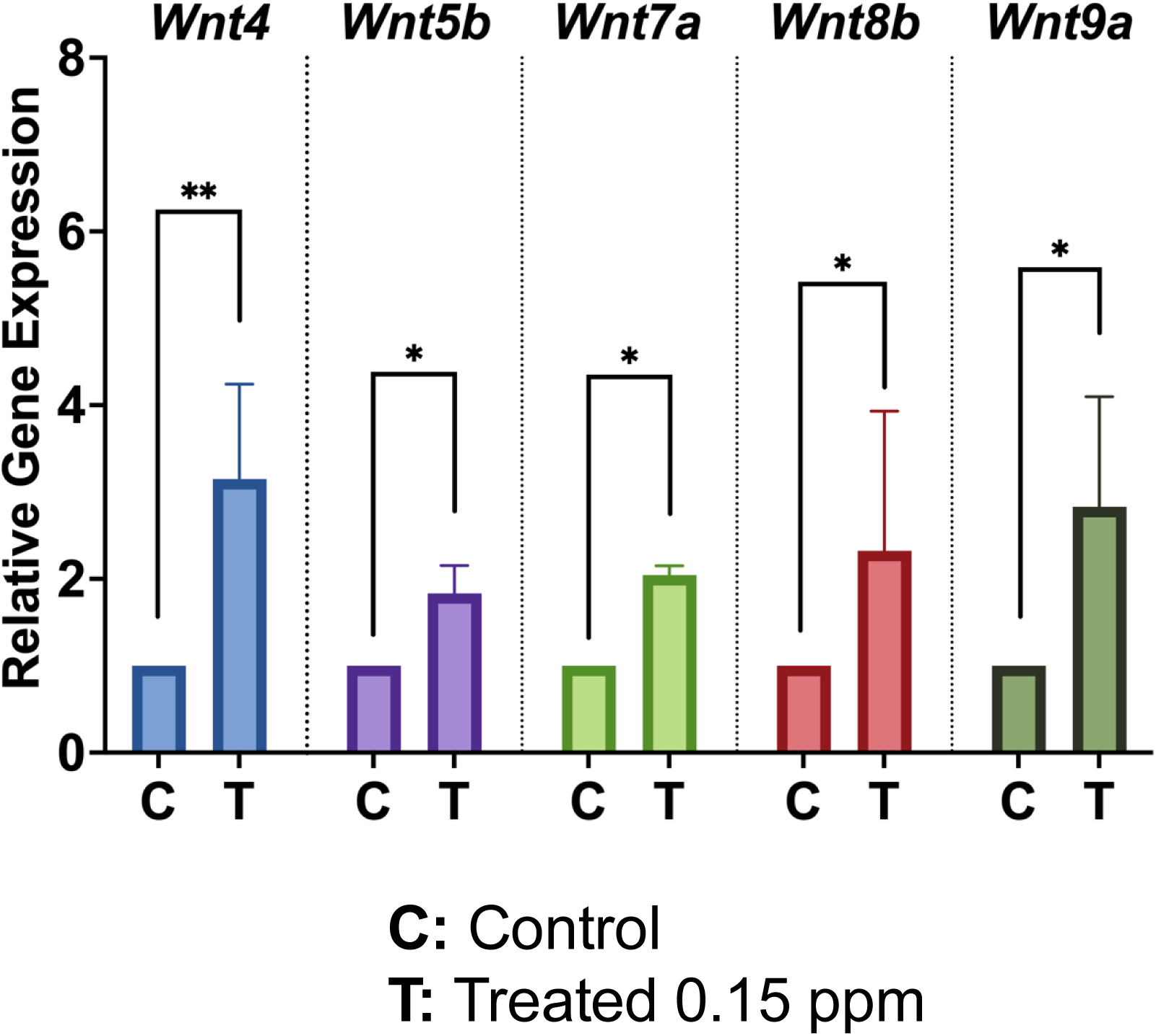
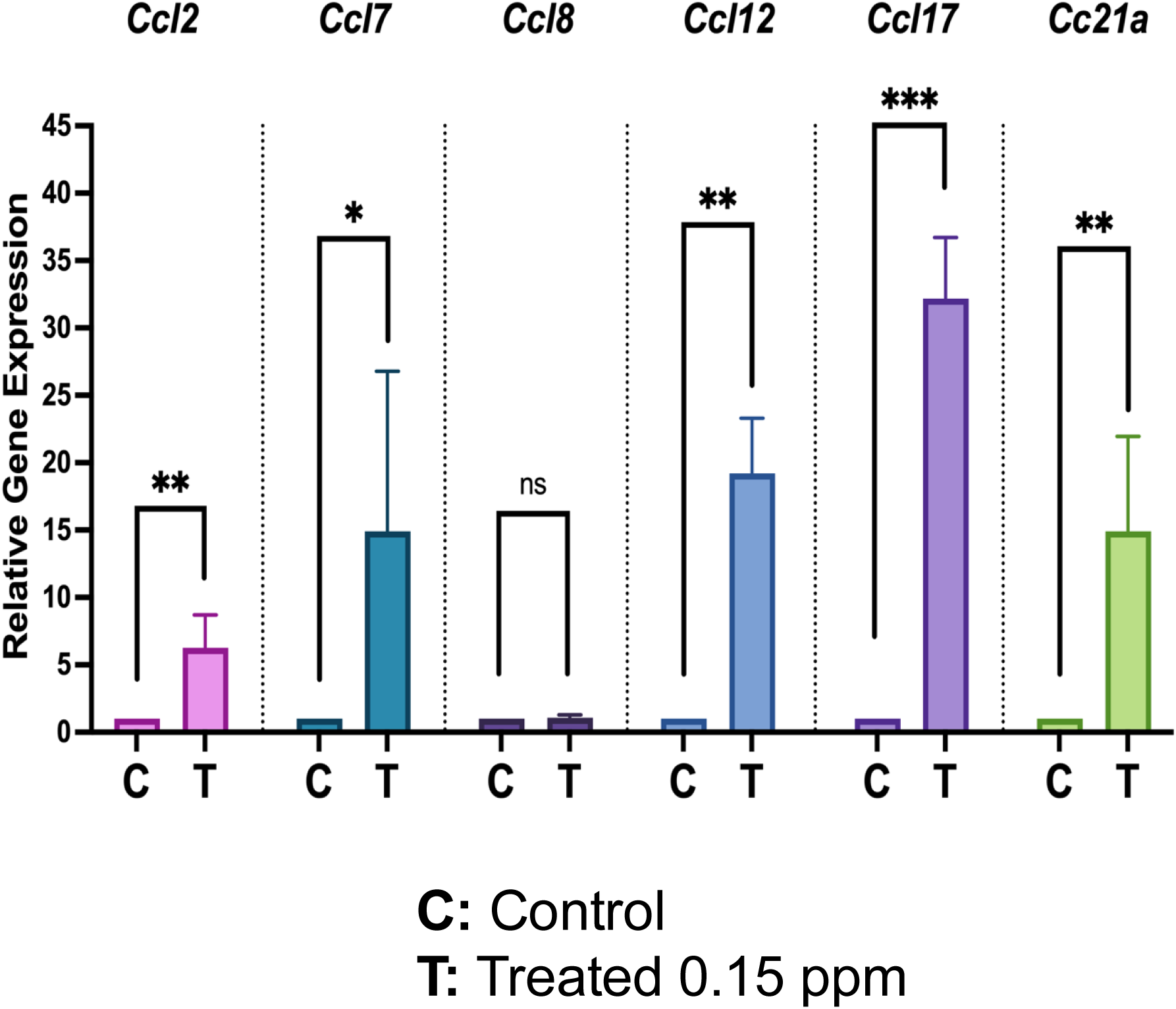
Exposure to a phthalate mixture leads to elevated expression of genes regulated by estrogen, Wnts, and chemokines in the uterus. Uteri collected at 6 months were used for quantitative real-time polymerase chain reaction (qPCR) analysis. Uterine RNA from control and 0.15 ppm groups were subjected to qPCR using primers specific for Hif2a, Cx43, Muc1, Fgf9, Ccn1, Pbx3, Six1, and Wt1 (**A**); primers specific for Wnt4, Wnt5b, Wnt7a, Wnt8a, and Wnt9b (**B**); primers specific for Ccl2, Ccl7, Ccl8, Ccl12, Ccl17 and Ccl21a (**C**). 36b4 was used as the internal control. Data represent mean ± SEM from six separate samples. Asterisks indicate statistically significant differences (*P < 0.1, * P < 0.05, ** P < 0.01).

We next investigated if phthalate-mediated increased estrogen-dependent signaling promotes epithelial proliferation to cause endometrial hyperplasia. Hence, we performed immunofluorescence (IF) staining for KI67, a well-known marker for cell proliferation, and found a dramatic increase in the number of KI67- positive cells in the uterine glandular epithelium (Fig. 3). In mice, endometrial hyperplasia is characterized by certain histopathological features. While the features can vary in severity, endometrial hyperplasia is generally accompanied by an increased number of endometrial glands that cause an altered gland to stroma ratio, and the stroma exhibits fibrotic changes with infiltration of immune cells.

**Figure 3:**
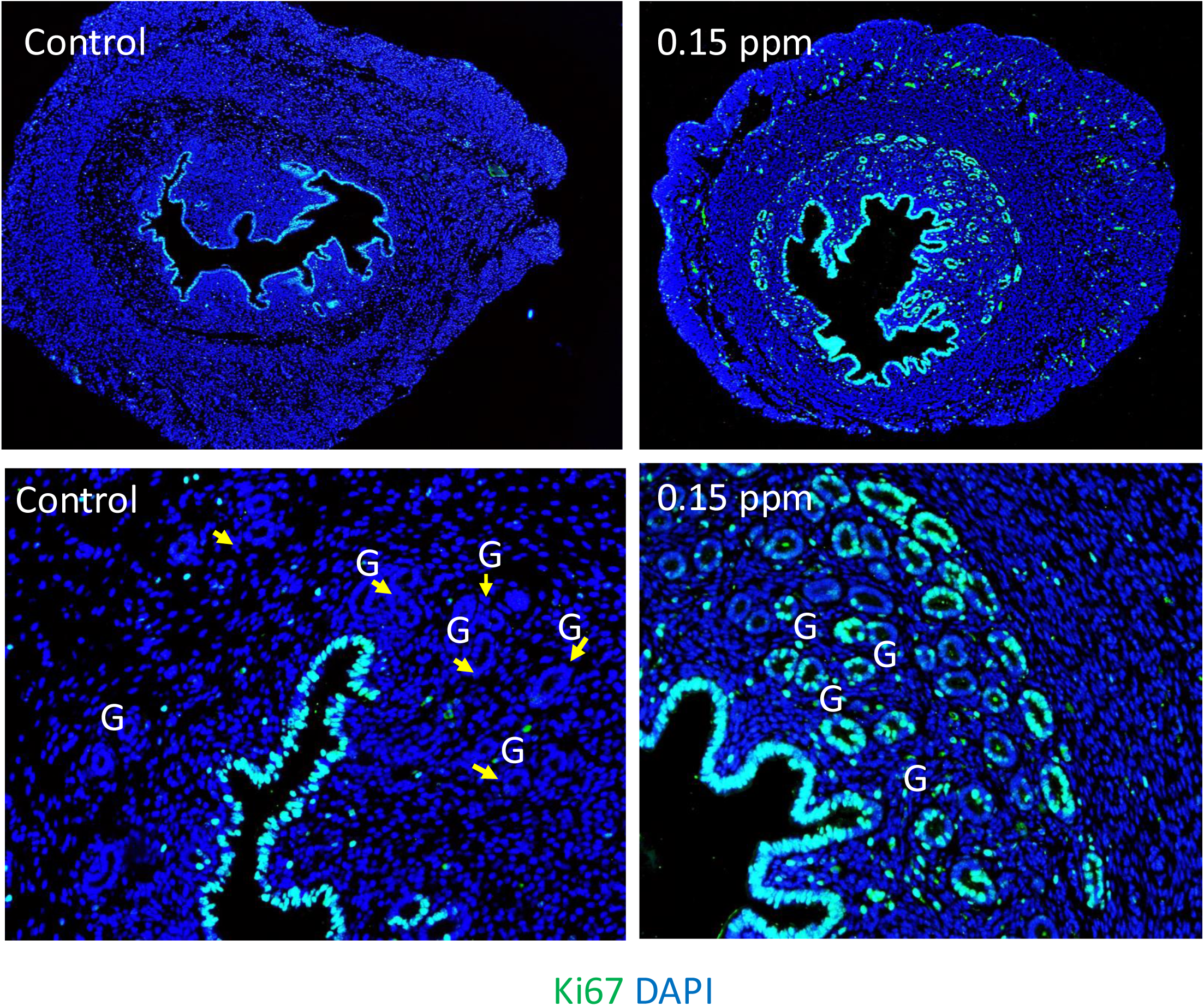
Enhanced proliferation of glandular epithelial cells in the uterus by long-term exposure to phthalates. Immunohistochemical localization of KI67 in the uterine sections of mice with or without exposure to 0.15 ppm phthalate mixture (N=3 per group). Upper panels indicate lower magnification (20x), and lower panels indicate higher magnification (40x). G indicates glands. The representative images from the control and phthalate mixture treated groups are shown.

We then evaluated stromal fibrosis in unexposed and phthalate-exposed uteri using Picrosirius red stain. The picrosirius red stain highlights the natural birefringence of collagen fibers under polarized light. Under this condition, the collagen is marked by a bright red color, and the intensity of the red color indicates the amount of collagen content. We noted that in the control group, the collagen fibers were barely visible in the uterus, but the phthalate-exposed samples exhibited a pronounced red color, indicating increased collagen deposition in the endometrium and myometrium (Fig. 4). These results indicated uterine fibrosis upon long-term exposure to a mixture of phthalates. To assess the infiltration of immune cells in the uterus, we next performed IF staining for F4/80, a specific marker for macrophages (Fig. 5). We observed a marked increase in the homing of macrophages in the endometrium in response to phthalate exposure. Interestingly, most of the macrophages were localized in the subepithelial stroma of phthalate-exposed uteri. This is consistent with the observation that blood vessels are beneath the epithelial cells in the nonpregnant mouse uterus (41). Taken together, these results indicated that the uterus exhibits features of hyperplasia upon long-term exposure to the phthalate mixture.

**Figure 4:**
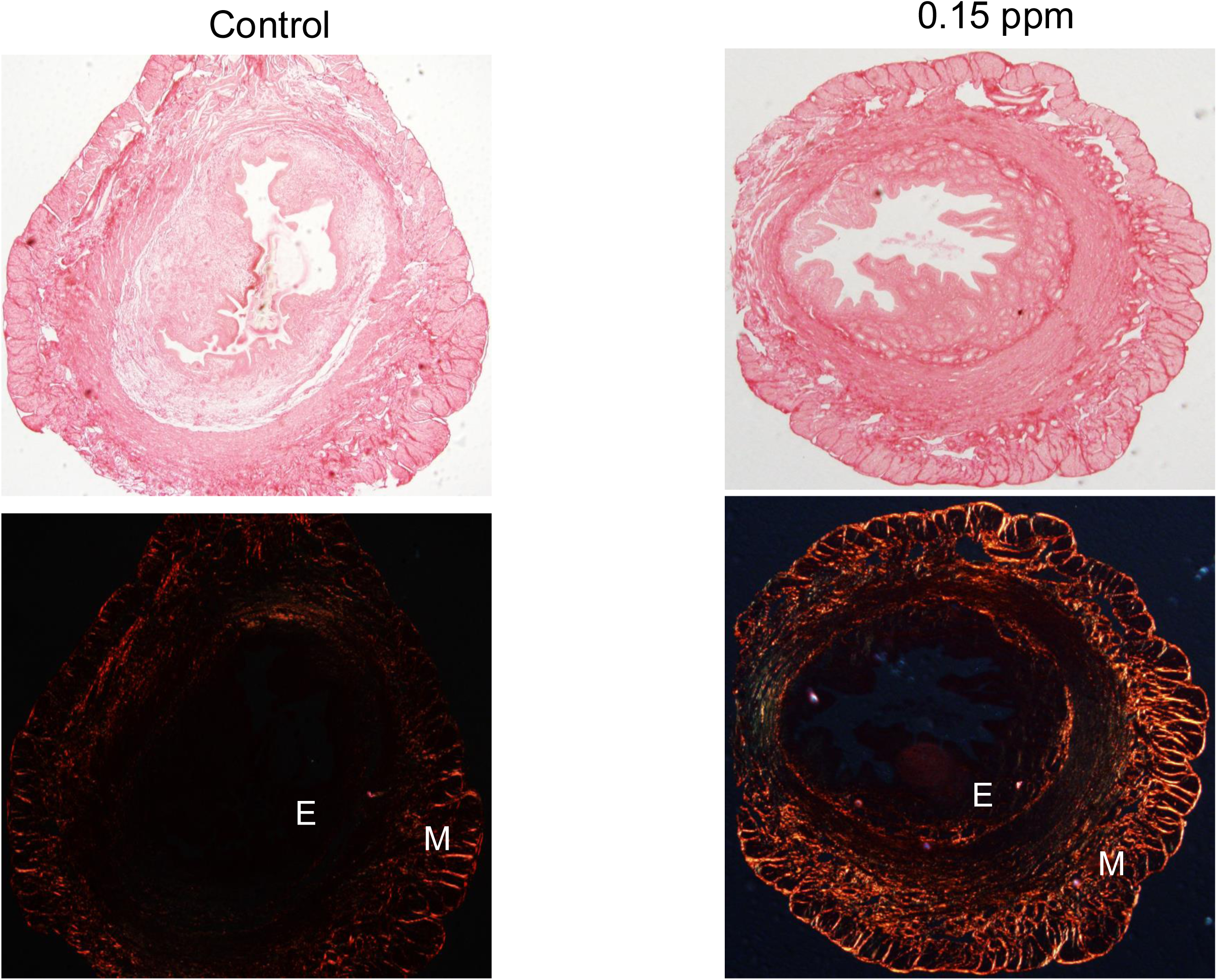
Exposure to a phthalate mixture leads to uterine fibrosis. Upper: Picro-Sirius red staining of uterine cross-sections of a control mouse (left) and 0.15 ppm group mouse (right). **Lower:** Visualization of collagen fibers in the upper panels under polarized light. N = 3 per group. E indicates endometrium, and M indicates myometrium. The representative images from the control and phthalate mixture treated groups are shown.

**Figure 5:**
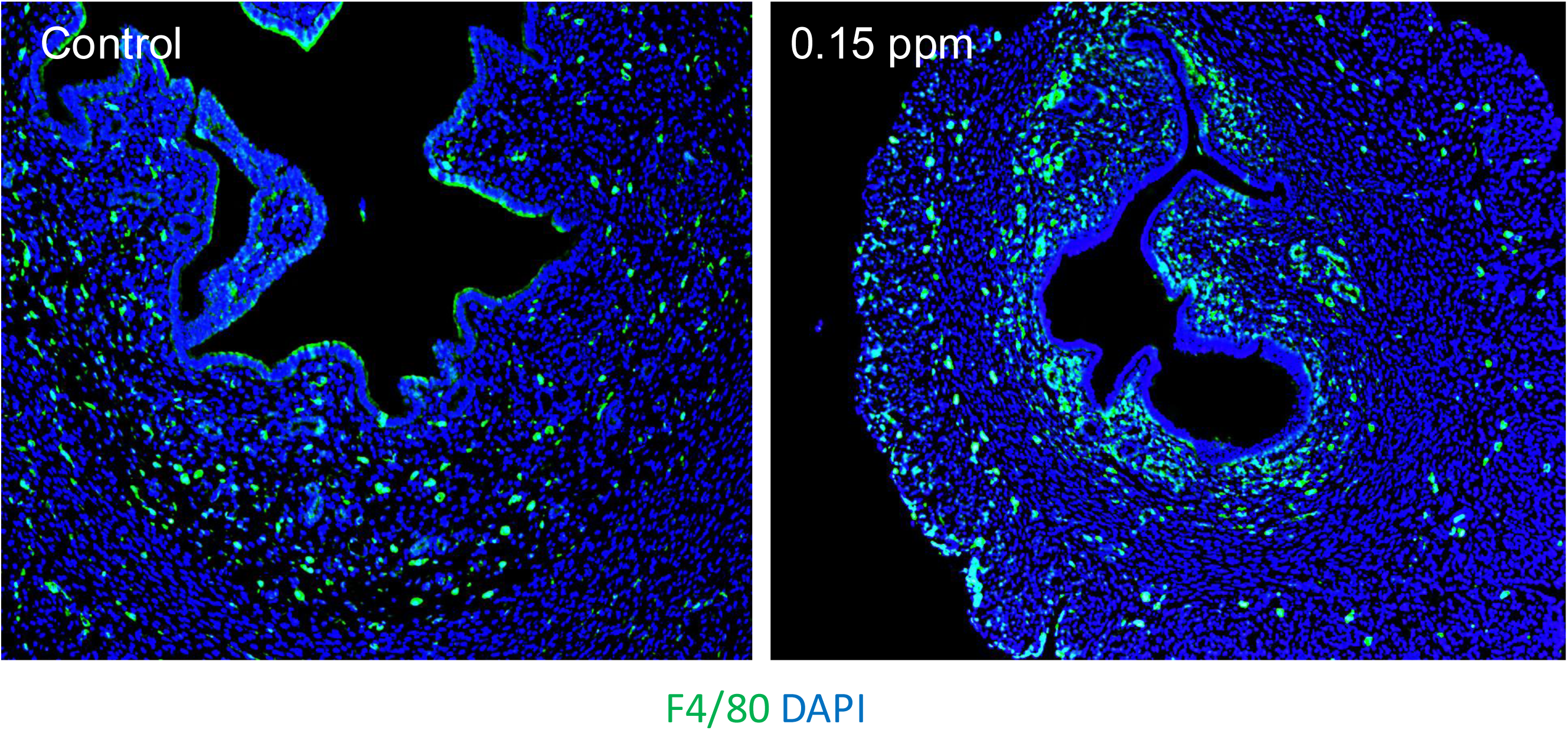
Exposure to the phthalate mixture promotes an influx of macrophages in the uterus. Uterine sections from control and 0.15 ppm treated groups were subjected to immunofluorescence using macrophage-specific F4/80 antibody. The representative images from the control and phthalate mixture treated groups are shown.

Several signaling pathways play a role in the development and progression of endometrial hyperplasia. The predominant signaling pathways activated in endometrial hyperplasia include the PI3K/AKT, MAPK/ERK, and Wnt/β-Catenin pathways (42–45). Each of these pathways promotes cell proliferation, leading to the hyperproliferative state of the endometrial cells. To identify the pathway that mediates the phthalate mixture-induced enhanced proliferation of endometrial epithelial cells, we investigated the expression levels of the regulatory proteins involved in each pathway. Interestingly, we observed a marked elevation in phospho-β-Catenin expression (Fig. 6) while phospho-ERK and phospho-AKT remained unchanged (data not shown), indicating an upregulation of Wnt/β-Catenin pathway in phthalate-exposed uteri.

**Figure 6:**
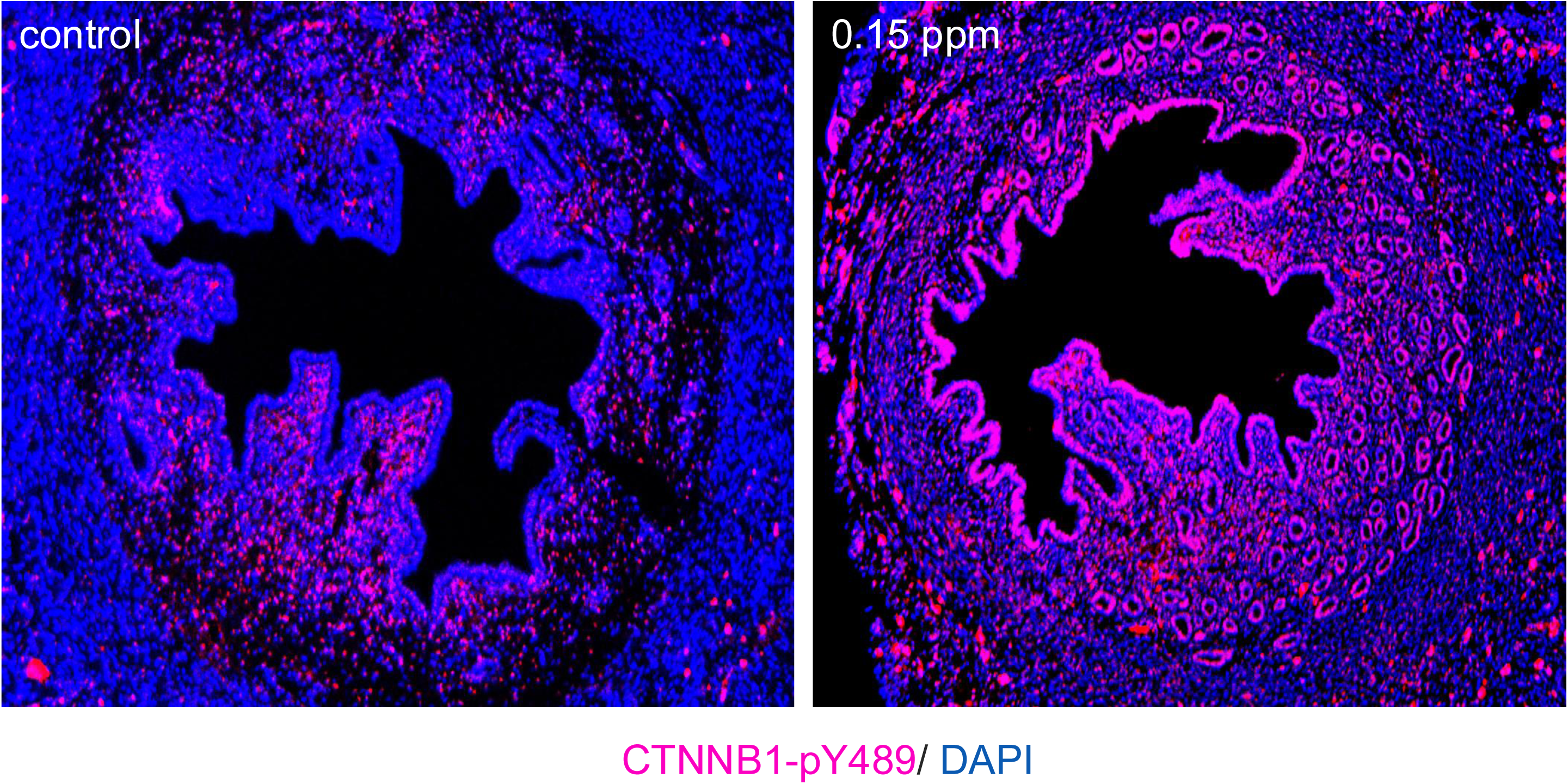
Activation of β-Catenin signaling pathway in the uterus by chronic phthalate exposure. Immunofluorescence analysis of phospho CTNNB1 in uterine sections of a control mouse (left) and 0.15 ppm group mouse (right). The representative images from the control and phthalate mixture treated groups are shown.

## DISCUSSION

Phthalate exposure in the general population is ubiquitous, and humans are repeatedly and continually exposed to phthalates (10–13). Studies indicate that phthalate metabolites are present in nearly 100% of tested human urine samples (10–13). These observations highlight the importance of understanding adverse health outcomes associated with phthalate exposure. Previous research indicated that prenatal exposure to a phthalate mixture results in multigenerational and transgenerational effects on uterine morphology in mice (46). However, humans experience daily phthalate exposure throughout their lives. In this study, six-week-old mice were continuously administered the phthalate mixture for 1 year. Moreover, the phthalate exposure occurred via food chow, mimicking the route of human exposure. Our results show that chronic exposure to a phthalate mixture comparable to what the EPA deems safe for human exposure can lead to undesirable uterine pathologies such as endometrial hyperplasia.

Aberrant proliferation is a hallmark of hyperplasia, a condition characterized by an increase in cell number without changes to cell morphology. Hyperplasia is not itself cancer but can develop into cancer with further genetic mutations or environmental insults (47). Here, we report that exposure to a mixture of phthalates leads to aberrant proliferation of the uterine epithelium, as indicated by KI67 staining. Notably, proliferation was particularly robust in the epithelium of the uterine glands, which are predominantly the site of origin for endometrial hyperplasia. Previous studies have examined the association between phthalate exposure and uterine conditions, including endometriosis and fibroids (48,49). Our study shows that chronic exposure to a mixture of phthalates in adults can have deleterious effects on the uterine glands, causing an increased gland:stroma ratio suggestive of endometrial hyperplasia. The impact on the uterine glands is pronounced at a low dose. This is not surprising since endocrine-disrupting chemicals such as phthalates have been shown to exhibit nonmonotonic dose responses (50). In light of these observations, it is concerning that exposure to a low concentration of phthalates, which may not be readily detectable in humans, is causing endometrial hyperplasia.

We report here that chronic phthalate exposure causes an upregulation of estrogen-regulated genes, many of which are involved in epithelial cell proliferation in uterine tissue (35–40). The significance of this finding is underscored by the fact that endometrial hyperplasia is often caused by prolonged exposure to estrogen without adequate progesterone to balance estrogen action (51). Prominent estrogen-regulated genes elevated by phthalate exposure are *Muc1*, *Fgf9*, *Ccn1*, *Cx43*, and *Hif2α*. Mucin 1, or Muc1, is a glycoprotein expressed on the uterine epithelium, and overexpression of Muc1 has been linked to endometrial cancers (52,53). Fgf9 promotes cell proliferation and survival in various tissues (54). In the endometrium, elevated Fgf9 expression might contribute to excessive cellular proliferation, leading to endometrial hyperplasia. Ccn1 is part of the Ccn family of matricellular proteins, which play a crucial role in various cellular processes, including cell proliferation, adhesion, migration, and angiogenesis (55). In endometrial cancer, overexpression of Ccn1 is linked to enhanced tumor growth, angiogenesis, and metastasis (55). Connexin 43 (Cx43), is a gap junction protein that plays a critical role in cell-to-cell communication by forming channels that allow the exchange of ions, metabolites, and signaling molecules between adjacent cells. The role of Cx43 in endometrial cancer is not known. Hypoxia-inducible factor 2 alpha or Hif2α is a transcription factor that plays a crucial role in the cellular response to hypoxia, a common feature in rapidly growing tumors, including endometrial cancer (56). Thus, dysregulated estrogen-dependent genes by long-term phthalate exposure drive the hallmarks of endometrial hyperplasia and underscore the importance of phthalate exposure on uterine homeostasis.

Our study also revealed an elevation in the expression of Pbx3, Six1, and Wt1, commonly overexpressed in various epithelial adenocarcinomas, including endometrial cancer. Pbx3 is a transcription factor that plays a critical role in the pathogenesis of endometrial cancer by promoting cancer cell proliferation, migration, and invasion (57). Six1 is a homeobox transcription factor involved in embryonic development and organogenesis (58). Six1 is known to regulate epithelial proliferation and is overexpressed in endometrial carcinoma. It plays a significant role in endometrial cancer by driving processes such as Epithelial-Mesenchymal Transition (EMT) and tumor progression (58). Wilms’ Tumor 1, or Wt1, is a gene that plays a role in the development of certain types of cancer, including endometrial cancer. It’s a transcription factor that is typically involved in the regulation of cell growth and differentiation. In endometrial cancer, WT1 can be overexpressed or mutated, which may contribute to tumorigenesis (59).

While the RNA-seq analysis revealed a significant upregulation of these genes by 6 months of phthalate exposure, the endometrial hyperplasia did not manifest until 1 year. At 6 months, an in-depth examination of uterine histology revealed an onset of fibrosis upon chronic exposure of mice to phthalates (28). Previous studies have shown that activated fibroblasts in fibrotic tissues secrete growth factors, extracellular matrix (ECM) proteins, and proteases that remodel the microenvironment and promote the proliferation of nearby epithelial cells through paracrine signaling (60). Further, persistent inflammation is a hallmark of fibrosis and a well-known risk factor for cancer. Activated immune cells release cytokines and chemokines, potentially playing a crucial role in the development of endometrial hyperplasia (40). Consistent with this notion, we found a marked upregulation of several chemokines, including Ccl12, Ccl17, Ccl2, Ccl21a, and Ccl7 in the uterus upon long-term treatment of phthalates. The influx of macrophages was also significantly elevated in the uterus in response to phthalate exposure. Future studies will address whether the measurement of inflammatory cytokines could serve as biomarkers for inflammation associated with phthalate exposure and endometrial hyperplasia.

An important finding of this study was the dramatic activation of the β-catenin pathway when mice were exposed long-term to a mixture of phthalates. Several Wnt ligands are induced in the uterus in response to phthalate exposure. Notably, previous studies have shown that Wnt4 is regulated by estrogen in the uterus, and it is upregulated in endometrial cancer (61,62). Upon activation of Wnt signaling, β-catenin accumulates in the nucleus and initiates the transcription of genes that promote cell proliferation. Indeed, aberrant activation of this pathway leads to uncontrolled cell proliferation and plays a significant role in the development and progression of endometrial cancer (44). It has been previously reported that a genetic mouse model with stabilized β-catenin expression displays endometrial glandular hyperplasia and not cancer, indicating that alteration in the β-catenin pathway is not sufficient for the development of endometrial cancer (63). Interestingly, a recent study has shown that developmental Diethylstilbestrol (DES) exposure in mice causes adenocarcinoma by activating Wnt/β-catenin and PI3K/AKT signaling pathways (64). These findings are consistent with our results, which show glandular hyperplasia in the phthalate-exposed mouse model displaying upregulation of Wnt/β-catenin and not PI3K/AKT signaling pathway.

In summary, chronic exposure to the phthalate mixture results in the dysregulation of estrogen and Wnt/β- Catenin signaling pathways, ultimately leading to the development of endometrial hyperplasia. Activation of these pathways contributes to abnormal glandular cell proliferation, tissue remodeling, and inflammation in the uterus. Future studies will address how estrogen and β-Catenin signaling pathways converge to promote endometrial hyperplasia in phthalate-exposed uteri. Insights into these signaling pathways highlight the potential mechanism by which phthalates induce endometrial hyperplasia and underscore the significance of understanding environmental factors in the pathogenesis of endometrial disorders.

## Acknowledgments

This work was supported by NIH (R01 ES032163, R01 ES034112, and T32 ES007326).

## REFERENCES

1. Pagoni A, Arvaniti OS, Kalantzi OI. Exposure to phthalates from personal care products: Urinary levels and predictors of exposure. Environ Res. 2022;212(Pt A):113194.

2. Carlos KS, de Jager LS, Begley TH. Determination of phthalate concentrations in paper-based fast food packaging available on the U.S. market. Food Addit Contam Part A Chem Anal Control Expo Risk Assess. 2021;38(3):501–512.

3. Gore AC, Chappell VA, Fenton SE, Flaws JA, Nadal A, Prins GS, Toppari J, Zoeller RT. Executive Summary to EDC-2: The Endocrine Society’s Second Scientific Statement on Endocrine-Disrupting Chemicals. Endocr Rev. 2015;36(6):593–602.

4. Gao X, Yang B, Tang Z, Luo X, Wang F, Xu H, Cai X. Determination of phthalates released from paper packaging materials by solid-phase extraction-high-performance liquid chromatography. J Chromatogr Sci. 2014;52(5):383–389.

5. Cao XL. Phthalate Esters in Foods: Sources, Occurrence, and Analytical Methods. Compr Rev Food Sci Food Saf. 2010;9(1):21–43.

6. Halden RU. Plastics and health risks. Annu Rev Public Health. 2010;31:179–194.

7. Zhang X, Chen Z. Observing phthalate leaching from plasticized polymer films at the molecular level. Langmuir. 2014;30(17):4933–4944.

8. Wittassek M, Koch HM, Angerer J, Bruning T. Assessing exposure to phthalates - the human biomonitoring approach. Mol Nutr Food Res. 2011;55(1):7–31.

9. Wu Y, Song Z, Little JC, Zhong M, Li H, Xu Y. An integrated exposure and pharmacokinetic modeling framework for assessing population-scale risks of phthalates and their substitutes. Environ Int. 2021;156:106748.

10. Silva MJ, Barr DB, Reidy JA, Malek NA, Hodge CC, Caudill SP, Brock JW, Needham LL, Calafat AM. Urinary levels of seven phthalate metabolites in the U.S. population from the National Health and Nutrition Examination Survey (NHANES) 1999-2000. Environmental health perspectives. 2004;112(3):331–338.

11. Hogberg J, Hanberg A, Berglund M, Skerfving S, Remberger M, Calafat AM, Filipsson AF, Jansson B, Johansson N, Appelgren M, Hakansson H. Phthalate diesters and their metabolites in human breast milk, blood or serum, and urine as biomarkers of exposure in vulnerable populations. Environmental health perspectives. 2008;116(3):334–339.

12. Heudorf U, Mersch-Sundermann V, Angerer J. Phthalates: toxicology and exposure. Int J Hyg Environ Health. 2007;210(5):623–634.

13. James-Todd T, Stahlhut R, Meeker JD, Powell SG, Hauser R, Huang T, Rich-Edwards J. Urinary phthalate metabolite concentrations and diabetes among women in the National Health and Nutrition Examination Survey (NHANES) 2001-2008. Environmental health perspectives. 2012;120(9):1307–1313.

14. Kay VR, Chambers C, Foster WG. Reproductive and developmental effects of phthalate diesters in females. Crit Rev Toxicol. 2013;43(3):200–219.

15. Zhang Y, Lin L, Cao Y, Chen B, Zheng L, Ge RS. Phthalate levels and low birth weight: a nested case-control study of Chinese newborns. J Pediatr. 2009;155(4):500–504.

16. Zhao Y, Chen L, Li LX, Xie CM, Li D, Shi HJ, Zhang YH. Gender-specific relationship between prenatal exposure to phthalates and intrauterine growth restriction. Pediatric research. 2014;76(4):401–408.

17. Factor-Litvak P, Insel B, Calafat AM, Liu X, Perera F, Rauh VA, Whyatt RM. Persistent Associations between Maternal Prenatal Exposure to Phthalates on Child IQ at Age 7 Years. PLoS One. 2014;9(12):e114003.

18. Toft G, Jonsson BA, Lindh CH, Jensen TK, Hjollund NH, Vested A, Bonde JP. Association between pregnancy loss and urinary phthalate levels around the time of conception. Environmental health perspectives. 2012;120(3):458–463.

19. Ferguson KK, McElrath TF, Ko YA, Mukherjee B, Meeker JD. Variability in urinary phthalate metabolite levels across pregnancy and sensitive windows of exposure for the risk of preterm birth. Environ Int. 2014;70:118–124.

20. Brehm E, Rattan S, Gao L, Flaws JA. Prenatal Exposure to Di(2-Ethylhexyl) Phthalate Causes Long-Term Transgenerational Effects on Female Reproduction in Mice. Endocrinology. 2018;159(2):795–809.

21. Schmidt JS, Schaedlich K, Fiandanese N, Pocar P, Fischer B. Effects of di(2-ethylhexyl) phthalate (DEHP) on female fertility and adipogenesis in C3H/N mice. Environmental health perspectives. 2012;120(8):1123–1129.

22. Richardson KA, Hannon PR, Johnson-Walker YJ, Myint MS, Flaws JA, Nowak RA. Di (2- ethylhexyl) phthalate (DEHP) alters proliferation and uterine gland numbers in the uteri of adult exposed mice. Reprod Toxicol. 2018;77:70–79.

23. Chiang C, Flaws JA. Subchronic Exposure to Di(2-ethylhexyl) Phthalate and Diisononyl Phthalate During Adulthood Has Immediate and Long-Term Reproductive Consequences in Female Mice. Toxicol Sci. 2019;168(2):620–631.

24. Chiang C, Lewis LR, Borkowski G, Flaws JA. Exposure to di(2-ethylhexyl) phthalate and diisononyl phthalate during adulthood disrupts hormones and ovarian folliculogenesis throughout the prime reproductive life of the mouse. Toxicology and applied pharmacology. 2020;393:114952.

25. Kannan A, Davila J, Gao L, Rattan S, Flaws JA, Bagchi MK, Bagchi IC. Maternal high-fat diet during pregnancy with concurrent phthalate exposure leads to abnormal placentation. Sci Rep. 2021;11(1):16602.

26. Yazdy MM, Coull BA, Gardiner JC, Aguiar A, Calafat AM, Xiaoyun Y, Schantz SL, Korrick SA. A possible approach to improving the reproducibility of urinary concentrations of phthalate metabolites and phenols during pregnancy. J Expo Sci Environ Epidemiol. 2018;28(5):448–460.

27. Laws MJ, Meling DD, Deviney ARK, Santacruz-Marquez R, Flaws JA. Long-term exposure to di(2-ethylhexyl) phthalate, diisononyl phthalate, and a mixture of phthalates alters estrous cyclicity and/or impairs gestational index and birth rate in mice. Toxicol Sci. 2023;193(1):48–61.

28. Shukla R, Arshee MR, Laws MJ, Flaws JA, Bagchi MK, Wagoner Johnson AJ, Bagchi IC. Chronic exposure of mice to phthalates enhances TGF beta signaling and promotes uterine fibrosis. Reprod Toxicol. 2023;122:108491.

29. Neier K, Cheatham D, Bedrosian LD, Gregg BE, Song PXK, Dolinoy DC. Longitudinal Metabolic Impacts of Perinatal Exposure to Phthalates and Phthalate Mixtures in Mice. Endocrinology. 2019;160(7):1613–1630.

30. Neier K, Cheatham D, Bedrosian LD, Dolinoy DC. Perinatal exposures to phthalates and phthalate mixtures result in sex-specific effects on body weight, organ weights and intracisternal A-particle (IAP) DNA methylation in weanling mice. J Dev Orig Health Dis. 2019;10(2):176–187.

31. Neier K, Montrose L, Chen K, Malloy MA, Jones TR, Svoboda LK, Harris C, Song PXK, Pennathur S, Sartor MA, Dolinoy DC. Short- and long-term effects of perinatal phthalate exposures on metabolic pathways in the mouse liver. Environ Epigenet. 2020;6(1):dvaa017.

32. Kavlock R, Boekelheide K, Chapin R, Cunningham M, Faustman E, Foster P, Golub M, Henderson R, Hinberg I, Little R, Seed J, Shea K, Tabacova S, Tyl R, Williams P, Zacharewski T. NTP Center for the Evaluation of Risks to Human Reproduction: phthalates expert panel report on the reproductive and developmental toxicity of di(2-ethylhexyl) phthalate. Reprod Toxicol. 2002;16(5):529–653.

33. Koo HJ, Lee BM. Human monitoring of phthalates and risk assessment. J Toxicol Environ Health A. 2005;68(16):1379–1392.

34. Davila J, Laws MJ, Kannan A, Li Q, Taylor RN, Bagchi MK, Bagchi IC. Rac1 Regulates Endometrial Secretory Function to Control Placental Development. Plos Genet. 2015;11(8):e1005458.

35. Hewitt SC, Deroo BJ, Hansen K, Collins J, Grissom S, Afshari CA, Korach KS. Estrogen receptor-dependent genomic responses in the uterus mirror the biphasic physiological response to estrogen. Molecular endocrinology. 2003;17(10):2070–2083.

36. Vasquez YM, Nandu TS, Kelleher AM, Ramos EI, Gadad SS, Kraus WL. Genome-wide analysis and functional prediction of the estrogen-regulated transcriptional response in the mouse uterusdagger. Biol Reprod. 2020;102(2):327–338.

37. Laws MJ, Taylor RN, Sidell N, DeMayo FJ, Lydon JP, Gutstein DE, Bagchi MK, Bagchi IC. Gap junction communication between uterine stromal cells plays a critical role in pregnancy-associated neovascularization and embryo survival. Development. 2008;135(15):2659–2668.

38. Bhurke A, Kannan A, Neff A, Ma Q, Laws MJ, Taylor RN, Bagchi MK, Bagchi IC. A hypoxia-induced Rab pathway regulates embryo implantation by controlled trafficking of secretory granules. Proc Natl Acad Sci U S A. 2020;117(25):14532–14542.

39. Neff AM, Blanco SC, Flaws JA, Bagchi IC, Bagchi MK. Chronic Exposure of Mice to Bisphenol-A Alters Uterine Fibroblast Growth Factor Signaling and Leads to Aberrant Epithelial Proliferation. Endocrinology. 2019;160(5):1234–1246.

40. Zhao Y, Li Q, Katzenellenbogen BS, Lau LF, Taylor RN, Bagchi IC, Bagchi MK. Estrogen-induced CCN1 is critical for establishment of endometriosis-like lesions in mice. Molecular endocrinology. 2014;28(12):1934–1947.

41. Rockwell LC, Pillai S, Olson CE, Koos RD. Inhibition of vascular endothelial growth factor/vascular permeability factor action blocks estrogen-induced uterine edema and implantation in rodents. Biol Reprod. 2002;67(6):1804–1810.

42. Kim TH, Yoo JY, Kim HI, Gilbert J, Ku BJ, Li J, Mills GB, Broaddus RR, Lydon JP, Lim JM, Yoon HG, Jeong JW. Mig-6 suppresses endometrial cancer associated with Pten deficiency and ERK activation. Cancer Res. 2014;74(24):7371–7382.

43. Pavlidou A, Vlahos NF. Molecular alterations of PI3K/Akt/mTOR pathway: a therapeutic target in endometrial cancer. ScientificWorldJournal. 2014;2014:709736.

44. Fatima I, Barman S, Rai R, Thiel KWW, Chandra V. Targeting Wnt Signaling in Endometrial Cancer. Cancers (Basel). 2021;13(10).

45. McMellen A, Woodruff ER, Corr BR, Bitler BG, Moroney MR. Wnt Signaling in Gynecologic Malignancies. International journal of molecular sciences. 2020;21(12).

46. Li K, Liszka M, Zhou C, Brehm E, Flaws JA, Nowak RA. Prenatal exposure to a phthalate mixture leads to multigenerational and transgenerational effects on uterine morphology and function in mice. Reprod Toxicol. 2020;93:178–190.

47. Sanderson PA, Critchley HO, Williams AR, Arends MJ, Saunders PT. New concepts for an old problem: the diagnosis of endometrial hyperplasia. Human reproduction update. 2017;23(2):232–254.

48. Kim JH, Kim SH. Exposure to Phthalate Esters and the Risk of Endometriosis. Dev Reprod. 2020;24(2):71–78.

49. Iizuka T, Yin P, Zuberi A, Kujawa S, Coon JSt, Bjorvang RD, Damdimopoulou P, Pacyga DC, Strakovsky RS, Flaws JA, Bulun SE. Mono-(2-ethyl-5-hydroxyhexyl) phthalate promotes uterine leiomyoma cell survival through tryptophan-kynurenine-AHR pathway activation. Proc Natl Acad Sci U S A. 2022;119(47):e2208886119.

50. Inman Z, Flaws JA. Endocrine Disrupting Chemicals, Reproductive Aging, and Menopause: A Review. Reproduction. 2024.

51. Kim JJ, Kurita T, Bulun SE. Progesterone action in endometrial cancer, endometriosis, uterine fibroids, and breast cancer. Endocr Rev. 2013;34(1):130–162.

52. Morrison C, Merati K, Marsh WL, Jr., De Lott L, Cohn DE, Young G, Frankel WL. The mucin expression profile of endometrial carcinoma and correlation with clinical-pathologic parameters. Appl Immunohistochem Mol Morphol. 2007;15(4):426–431.

53. Engel BJ, Bowser JL, Broaddus RR, Carson DD. MUC1 stimulates EGFR expression and function in endometrial cancer. Oncotarget. 2016;7(22):32796–32809.

54. Turner N, Grose R. Fibroblast growth factor signalling: from development to cancer. Nat Rev Cancer. 2010;10(2):116–129.

55. Jia Q, Xu B, Zhang Y, Ali A, Liao X. CCN Family Proteins in Cancer: Insight Into Their Structures and Coordination Role in Tumor Microenvironment. Front Genet. 2021;12:649387.

56. Wicks EE, Semenza GL. Hypoxia-inducible factors: cancer progression and clinical translation. J Clin Invest. 2022;132(11).

57. Morgan R, Pandha HS. PBX3 in Cancer. Cancers (Basel). 2020;12(2).

58. Suen AA, Jefferson WN, Wood CE, Williams CJ. SIX1 Regulates Aberrant Endometrial Epithelial Cell Differentiation and Cancer Latency Following Developmental Estrogenic Chemical Exposure. Molecular cancer research : MCR. 2019;17(12):2369–2382.

59. McEachron J, Baqir AW, Zhou N, Jabbar A, Gupta R, Levitan D, Lee YC. Evaluation of the incidence and clinical significance of WT-1 expression in uterine serous carcinoma. Gynecol Oncol Rep. 2022;39:100918.

60. Caligiuri G, Tuveson DA. Activated fibroblasts in cancer: Perspectives and challenges. Cancer Cell. 2023;41(3):434–449.

61. Hou X, Tan Y, Li M, Dey SK, Das SK. Canonical Wnt signaling is critical to estrogen-mediated uterine growth. Molecular endocrinology. 2004;18(12):3035–3049.

62. Katayama S, Ashizawa K, Fukuhara T, Hiroyasu M, Tsuzuki Y, Tatemoto H, Nakada T, Nagai K. Differential expression patterns of Wnt and beta-catenin/TCF target genes in the uterus of immature female rats exposed to 17alpha-ethynyl estradiol. Toxicol Sci. 2006;91(2):419–430.

63. Jeong JW, Lee HS, Franco HL, Broaddus RR, Taketo MM, Tsai SY, Lydon JP, DeMayo FJ. beta-catenin mediates glandular formation and dysregulation of beta-catenin induces hyperplasia formation in the murine uterus. Oncogene. 2009;28(1):31–40.

64. Padilla-Banks E, Jefferson WN, Papas BN, Suen AA, Xu X, Carreon DV, Willson CJ, Quist EM, Williams CJ. Developmental estrogen exposure in mice disrupts uterine epithelial cell differentiation and causes adenocarcinoma via Wnt/beta-catenin and PI3K/AKT signaling. PLoS Biol. 2023;21(10):e3002334.

